# The developing retina undergoes mitochondrial remodeling via PINK1/PRKN-dependent mitophagy

**DOI:** 10.1101/2025.03.30.646079

**Authors:** Juan Zapata-Muñoz, Juan Ignacio Jiménez-Loygorri, Michael Stumpe, Beatriz Villarejo-Zori, Sandra Alonso-Gil, Petra Terešak, Benan J. Mathai, Ian G Ganley, Anne Simonsen, Jörn Dengjel, Patricia Boya

**Affiliations:** Department of Cellular and Molecular Biology, Centro de Investigaciones Biológicas Margarita Salas, CSIC, Madrid, Spain; Molecular Oncology Programme, Centro Nacional de Investigaciones Oncológicas (CNIO), Madrid, Spain; Dept of Molecular Cell Biology, Inst for Cancer Research, Oslo University Hospital; Centre for Cancer Cell Reprogramming, Institute of Clinical Medicine, Faculty of Medicine, University of Oslo, 1112 Blindern, Oslo 0317, Norway; MRC Protein Phosphorylation and Ubiquitylation Unit, University of Dundee, Dundee DD1 5EH, UK; Department of Biology, University of Fribourg, Fribourg, Switzerland; Department of Neuroscience and Movement Science, University of Fribourg, Chemin. du Musée 14, 1700 Fribourg, Switzerland

**Keywords:** mitophagy, autophagy, retina, development, PINK1

## Abstract

Mitophagy, the selective degradation of mitochondria, is essential for retinal ganglion cell (RGC) differentiation and retinal homeostasis. However, the specific mitophagy pathways involved and their temporal dynamics during retinal development and maturation remain poorly understood. Using proteomics analysis of isolated mouse retinas across developmental stages and the mitophagy reporter mouse line, *mito*-QC, we characterized mitophagy throughout retinogenesis. While mitolysosomes were more prevalent in the mature retina, we observed two distinct mitophagy peaks during embryonic development. The first, independent of PTEN-induced kinase 1 (PINK1) activation, was associated with RGCs. The second, PINK1-dependent peak was triggered after an increase in retinal oxidative stress. This PINK1-dependent, oxidative stress-induced mitophagy pathway is conserved in mice and zebrafish, providing the first evidence of programmed, PINK1-dependent mitophagy during development.

## Introduction

The retina is a complex organ comprising more than 60 types of neurons that process and transmit electrochemical signals to the brain, enabling vision [1]. Differentiation of retinal cells occurs in a highly orchestrated manner, spanning embryonic day (E) 8.5–9.0, when optic vesicles are formed in the mouse forebrain, to postnatal day (P) 14, when eye opening occurs [2]. Six main neuronal types are generated from the pool of retinal progenitor cells (RPCs): rod and cone photoreceptors, amacrine, bipolar, horizontal and retinal ganglion cells (RGCs); and Müller glia. This process is conserved across all vertebrates, RGCs being the first to differentiate at E11.5 in mice [3]. According to the competence model of differentiation, retinal development can be divided in two main stages: an embryonic period characterized by proliferation of multipotent RPCs [4], and a second postnatal period during which Müller glia differentiate and neurons mature and migrate to their final position within the trilaminar structure of the retina, where synaptogenesis occurs [5].

Autophagy is an intracellular degradation pathway through which cellular components are recycled within lysosomes. Mitophagy is the autophagic process by which mitochondria are selectively degraded and is critical for maintaining retinal homeostasis [6]. We have previously demonstrated a role for programmed mitophagy via the NIX-dependent pathway in RGC differentiation [7]. However, it remains unclear which developmental stages mitophagy is most involved in, and whether PINK1/PRKN-dependent mitophagy, more associated to stressful scenarios [8, 9], also contributes to retinal development.

Quantitative proteomics enables simultaneous identification and quantification of thousands of proteins, and is a crucial tool for studying complex biological systems. This approach has recently been employed to identify mitophagy proteins that drive the differentiation of induced neurons (iNeurons) *in vitro* [10]. While 27 human eye proteomic datasets have been published since 2013, only 7 correspond to the neuroretina and none address retinal development [11].

By combining proteomic analyses with mitophagy reporter animals during retinal development in mice and zebrafish, we have identified key shifts in the mitophagy profile at discrete stages of embryonic development. Moreover, we show that these changes are associated with distinct mitophagy pathways, including the stress-associated PINK1/PRKN-dependent mitophagy.

## Results

### Proteomic analysis reveals two distinct phases of mouse retinal development

We collected at least four replicates of isolated retinas at 9 different timepoints: embryonic stages E13.5, E14.5, E15.5, E16.5 and E18.5; and postnatal stages P0, P9, P15 and P30. These stages span from the first timepoint at which the retina can be easily separated from the retinal pigment epithelium (E13.5) to the formation of a fully mature and functional retina (P30) (**Fig. 1a**). To characterize proteome dynamics during retinogenesis, we performed a quantitative proteomics analysis using liquid chromatography–tandem mass spectrometry (LC-MS/MS) with data-independent acquisition (DIA) for peptide identification and quantification (**Fig. 1a**) (**Table S1**). Determining whether distinct developmental stages can be grouped based on their protein composition is essential to understand the process of retinal development and function. Unsupervised hierarchical clustering analysis (HCA) and principal component analysis (PCA) were used to group samples into two blocks: one corresponding to embryonic development (13.5–P0) and the other to postnatal retinal maturation (P9, P15 and P30) (**Fig. 1b**). Stage identity was validated using known retinal differentiation markers. Proteins essential for early retinal development and RPCs function, such as SOX9, RAX, PAX6, SIX3 and LHX2 [12], were highly abundant during early timepoints (**Fig. S1a**). Expression of proteins involved in horizontal, amacrine and bipolar cells [13], including TFAP2A, CALB1, CALB2, increased from E16.5 onwards (**Fig. S1a**). Levels of OTX2, which is involved in the early photoreceptor differentiation and bipolar cell survival and terminal differentiation [14], peaked at P9, and remained elevated in the mature retina (**Fig. S1a**), as confirmed by immunostaining (**Fig. S1b**). The remaining proteins, levels of which increased from E18.5 or P0 onwards, are involved in photoreceptor and Müller glia differentiation (**Fig. S1a**). The observed levels of these differentiation markers validate our proteomic data and corroborate previous studies of retinal development [15]. To explore the temporal dynamics of distinct proteome subsets, we performed an unsupervised k-means clustering, segregating proteins into nine clusters, and conducted over-representation analysis (ORA) using the KEGG pathways database (**Fig. 1c**). Cluster 2 (**Fig. 1c, orange**) was highly enriched at early embryonic and late post-natal stages, and contained terms related with cell cycle, DNA replication, and DNA repair. Proteins that form the spliceosome, the activity of which is implicated in differentiation processes [16], were found in Cluster 3 and, more robustly, in Cluster 7 (**Fig. 1c, yellow and light blue**). This supports the importance of alternative splicing in retinal development. Cluster 6 (Fig. 1c, dark green), enriched for synapse-associated terms, showed a marked decrease from P0 to P9, followed by recovery from P9 to P30, suggesting downregulation of synaptic proteins during synaptogenesis and subsequent upregulation in mature synapses. This cluster also included autophagy-related terms like “Phagosome,” “Lysosome,” “SNARE interactions,” and “mTOR signaling.” However, the “Autophagy” pathway itself was enriched in Cluster 4 (**Fig. 1c, light green**), highlighting its increasing importance during postnatal stages. Finally, Cluster 9 (**Fig. 1c, violet**) showed significant enrichment for “Mitophagy” and other mitochondrial metabolism-related terms, suggesting their prevalence after P0.

**Fig. 1.**
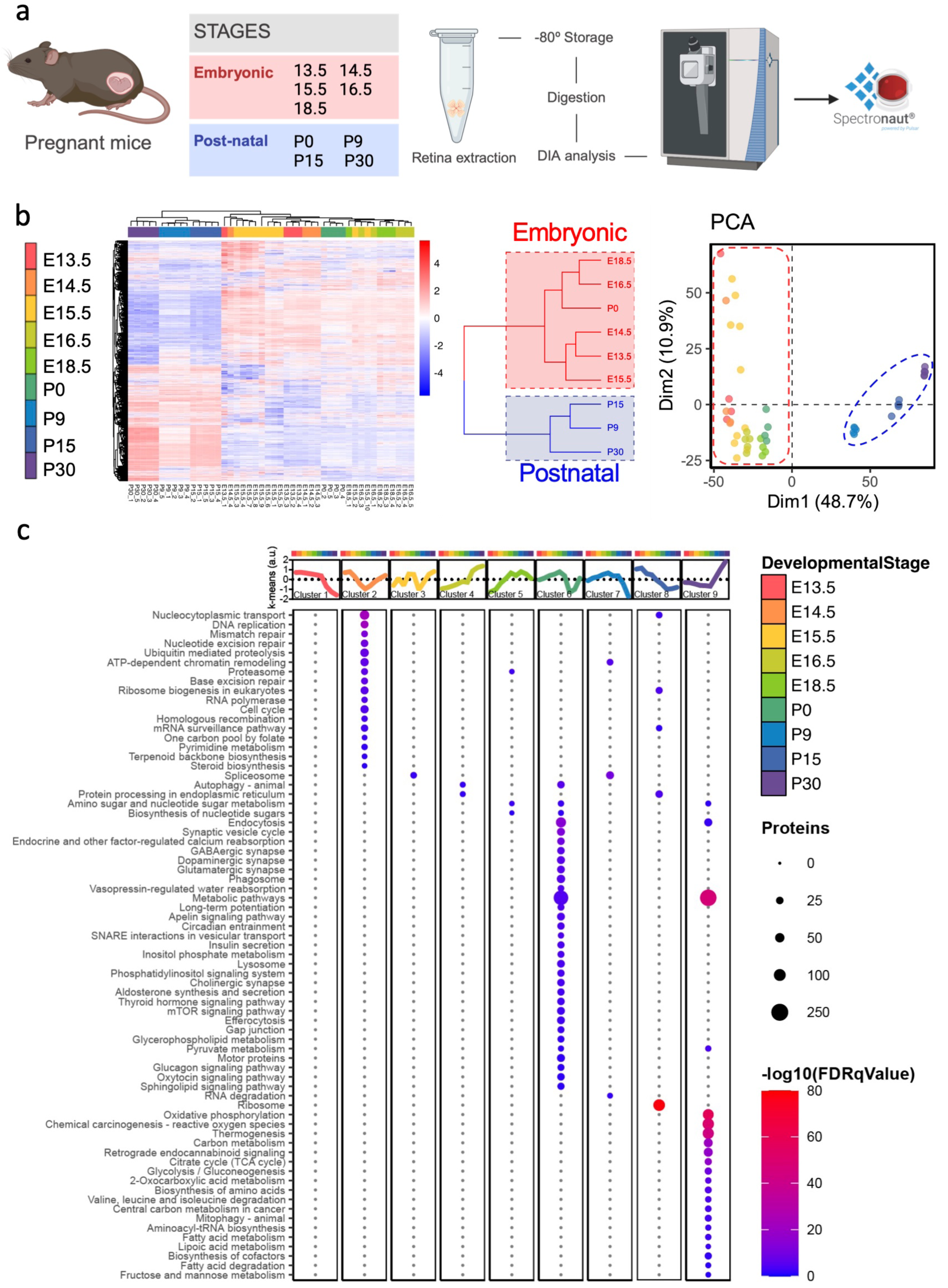
Temporal proteomics of the developing mouse retina. **(a)** Workflow for retinal sample acquisition and processing for LC-MS/MS data collection. Retinas were isolated and stored until all samples were obtained. Proteins were digested with trypsin, resulting peptides were extracted, and DIA-based quantitative LC-MS/MS was performed, followed by data analysis using Spectronaut software. **(b)** Hierarchical clustering and PCA of temporal proteomics data reveal two distinct developmental phases. **(c)** k-means clustering analysis of proteins grouped according to expression patterns. Circle size represents the number of proteins within the KEGG terms. Statistical significance is represented as -log10(FDR q-value). DIA, data-independent acquisition; FDR q-value, false discovery rate-adjusted q-value; KEGG, Kyoto Encyclopedia of Genes and Genomes; PCA, principal component analysis.

To further characterize these two developmental phases, we compared protein abundance between E13.5 and P0 (embryonic phase) and between P0 and P30 (postnatal phase) (Fig. S2a). Metabolic proteins (HK1, SLC16A1, and IDH3A) generally increased exponentially during the postnatal phase, while only the glycolytic enzyme HK1 also increased during embryonic development (Fig. S2b), supporting the importance of glycolysis in embryonic retinal stages, as previously demonstrated [7]. Conversely, asparagine synthetase (ASNS) levels increased during the embryonic phase but were extremely low postnatally (Fig. S2c), likely reflecting asparagine’s role in rapid cell division [17]. In line with this observation, NASP and RCC2, involved in DNA replication and repair, showed higher levels during the embryonic phase (Fig. S2c). Overall, our proteomic analysis revealed two distinct phases of retinal development: an embryonic phase characterized by proteins involved in cell division and spliceosome activity, and a postnatal phase marked by synapse-, metabolism-, and autophagy/mitophagy-related proteome remodeling.

### Mitochondrial biogenesis and mitophagy are upregulated in the postnatal retina

Given the over-representation of mitochondrial metabolism and mitophagy in postnatal stages (**Fig. 1c**), we performed a targeted analysis of mitochondrial proteins. Most mitochondrial proteins, including those of the electron transport chain and inner/outer and intermembrane compartments, increased after P0 (**Fig. 2a**). MTCO1 immunostaining confirmed this dynamic across different retinal stages (**Fig. 2b, c**). These findings indicate that expression of mitochondrial and mitophagy-related proteins (found in Cluster 9) is induced at the postnatal period. We next examined specific proteins involved in mitophagy. Proteins involved in LC3 lipidation, essential for autophagosome biogenesis [18], were upregulated during retinal development (**Fig. S3a**), except for ATG4B, which decreased over time (**Fig. S3a**), likely due to its role in autophagosome recycling [19]. Four mammalian ATG8/LC3 proteins (MAP1LC3A, MAP1LC3B, GABARAPL1, and GABARAPL2) detected in our proteomic pipeline also showed higher levels postnatally **(Fig. S3b**). Mitophagy receptors FUNDC1, PHB2, and BNIP3 increased after P0, while NIX remained stable, except for a marked decrease at P0 (**Fig. 3b**). Adaptor proteins recognizing polyubiquitin chains on damaged or stressed mitochondria [20] showed opposing patterns: TOLLIP and OPTN increased after birth, while SQSTM1 and NBR1 decreased during retinal development (**Fig. 3c**). The negative regulators of PINK1/PRKN- and NIX/BNIP3-dependent mitophagy, USP30 and PPTC7, respectively, showed slight fluctuations throughout retinal development (**Fig. 3d**). A slight downregulation of USP30 at E15.5 suggests potential PINK1/PRKN mitophagy activity at this stage. Both USP30 and PPTC7 increased from P0 to P30 (**Fig. 3d**), likely to limit both mitophagy pathways postnatally. While the autophagy core machinery clearly increases at the protein level, the interplay between different mitophagy pathways and their relative importance during retinal development requires further investigation.

**Fig. 2.**
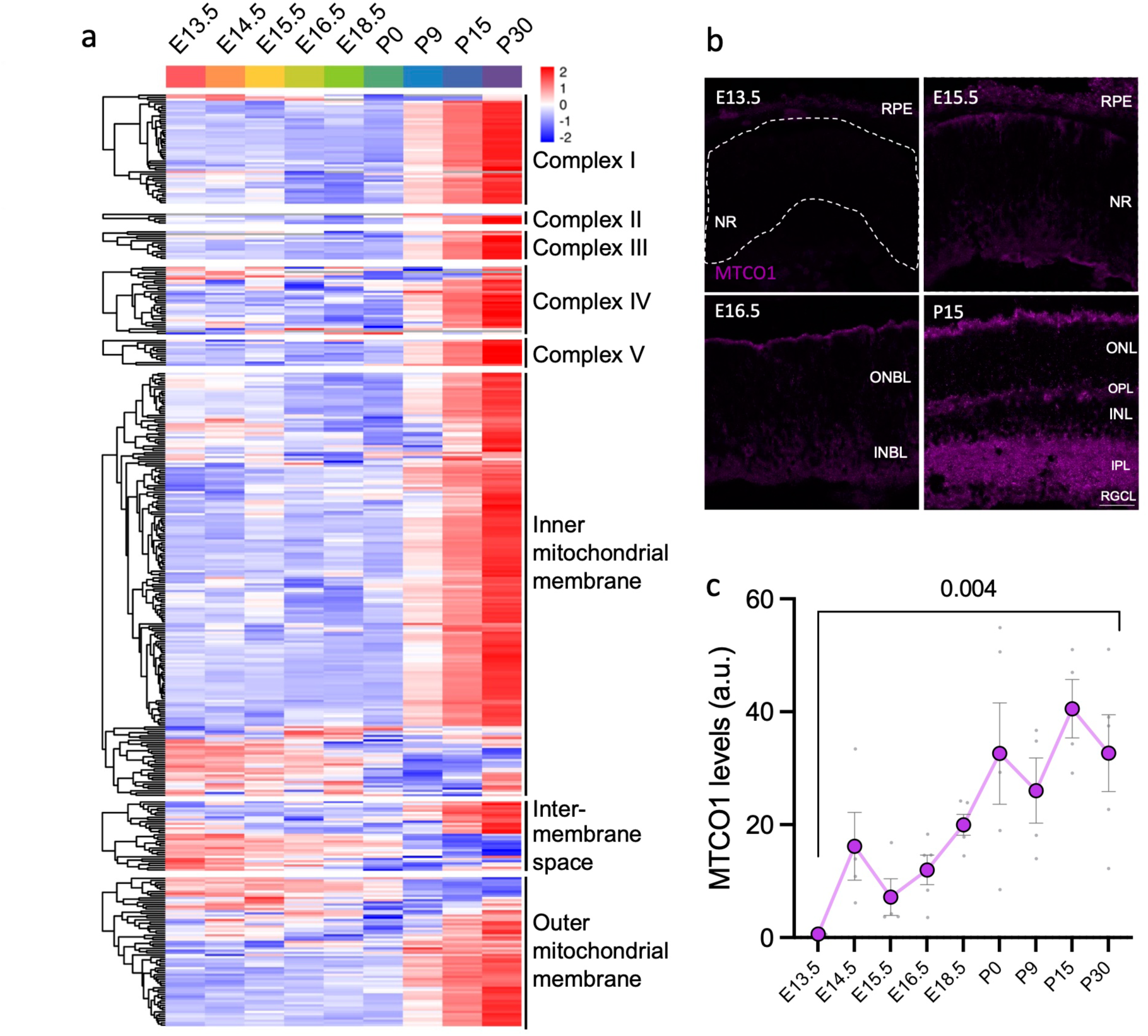
Expression of mitochondrial and OXPHOS-related proteins in the mouse retina increases after birth. **(a)** Heatmaps of mitochondrial protein expression during retinal development. **(b)** Representative images of central retinal cryosections from embryonic day (E)13.5, E15.5, E16.5, and postnatal day (P)15, immunolabeled for MTCO1 (purple). **(c)** Quantification of MTCO1-positive area during the different retinal developmental stages. Large dots represent the mean, and small dots represent individual mice (n = 2–5). Data are presented as mean ± SEM. *P* values were calculated using a two-tailed Student’s *t*-test. Scale bar: 40 μm. NR, neural retina; INBL, inner neuroblastic layer; INL, inner nuclear layer; IPL, inner plexiform layer; MTCO1, cytochrome c oxidase 1; ONBL, outer neuroblastic layer; ONL, outer nuclear layer; OPL, outer plexiform layer; OXPHOS, oxidative phosphorylation; RGCL, retinal ganglion cell layer; RPE, retinal pigment epithelium; SEM, standard error of the mean.

**Fig. 3.**
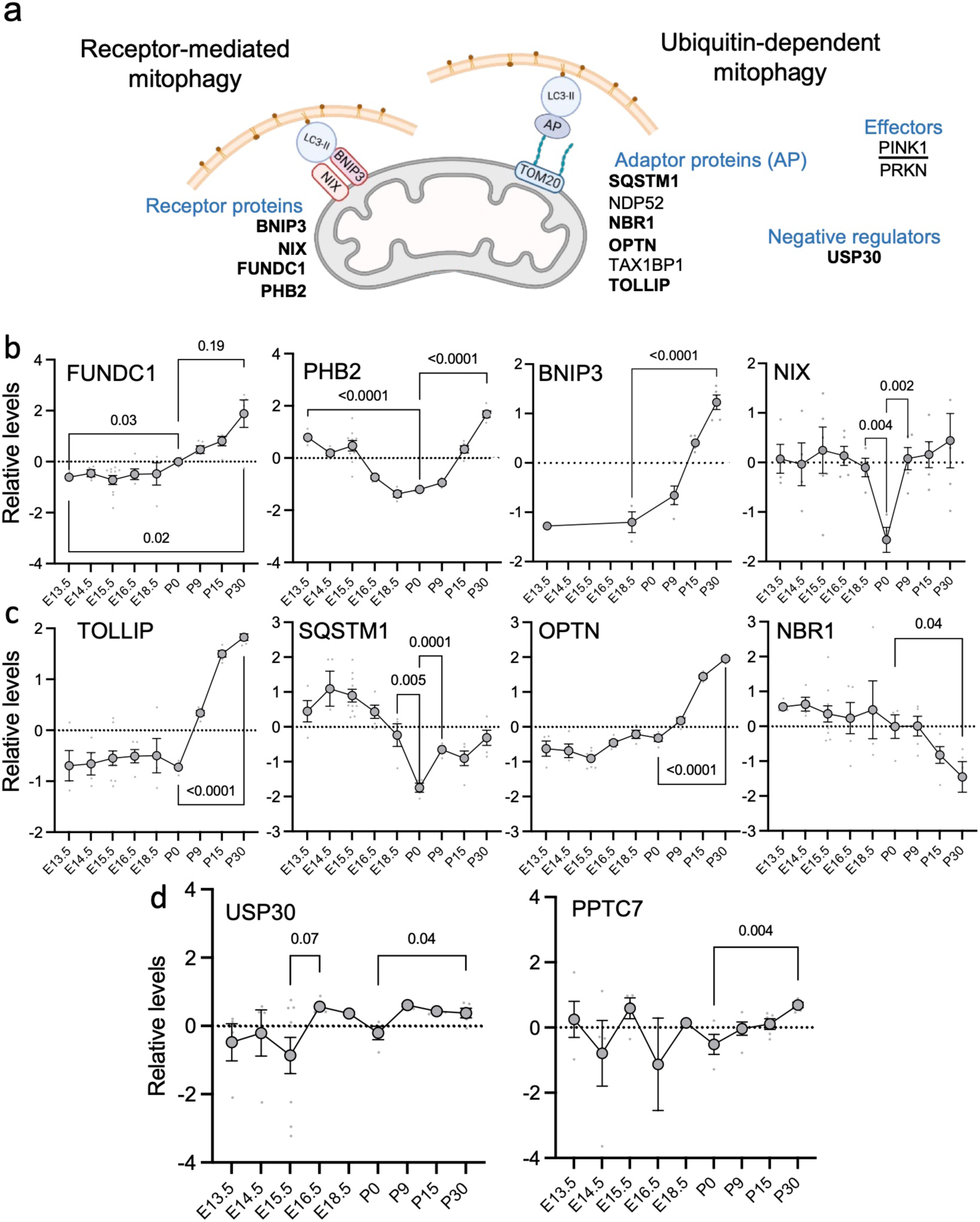
Mitophagy-related proteins exhibit distinct expression patterns during retinal development. **(a)** Diagram of mitophagy pathways. Relevant proteins are indicated, with those detected in the proteomic analysis highlighted. **(b–d)** Z-score normalized protein expression levels of mitophagy receptors (b), adaptor proteins (c), and PINK1/PRKN and NIX/BNIP3 negative regulators (d), detected by proteomics analysis. Large dots represent the mean, small gray dots represent individual mice (n ≥ 4), and bars represent the SEM. *P* values were calculated using a two-tailed Student’s *t*-test (PHB2, BNIP3, NIX; c and d) or a two-tailed Mann–Whitney *U*-test (FUNDC1). E, embryonic day; P, postnatal day; SEM, standard error of the mean.

We next used the mitophagy reporter mouse line *mito*-QC to quantify mitolysosomes at 9 developmental stages, from E10.5 to P30 (**Fig. 4a**). This reporter, comprising an mCherry-GFP tag fused to the FIS1 outer mitochondrial membrane localization signal, allows visualization of mitolysosomes as red puncta due to GFP quenching in the acidic lysosomal lumen [21]. Consistent with our proteomic data, mitophagy was most active in the mature neuroretina at P30 (**Fig. 4b-d**), with mitolysosomes concentrated in the outer nuclear layer (ONL), where photoreceptors reside, as previously reported [22]. However, two mitophagy peaks were also observed during embryonic retinal development, at E12.5–E13.5 and E16.5 (**Fig. 4b-d**), suggesting that mitophagy modulation during embryonic stages may be important for proper retinal cell type differentiation.

**Fig. 4.**
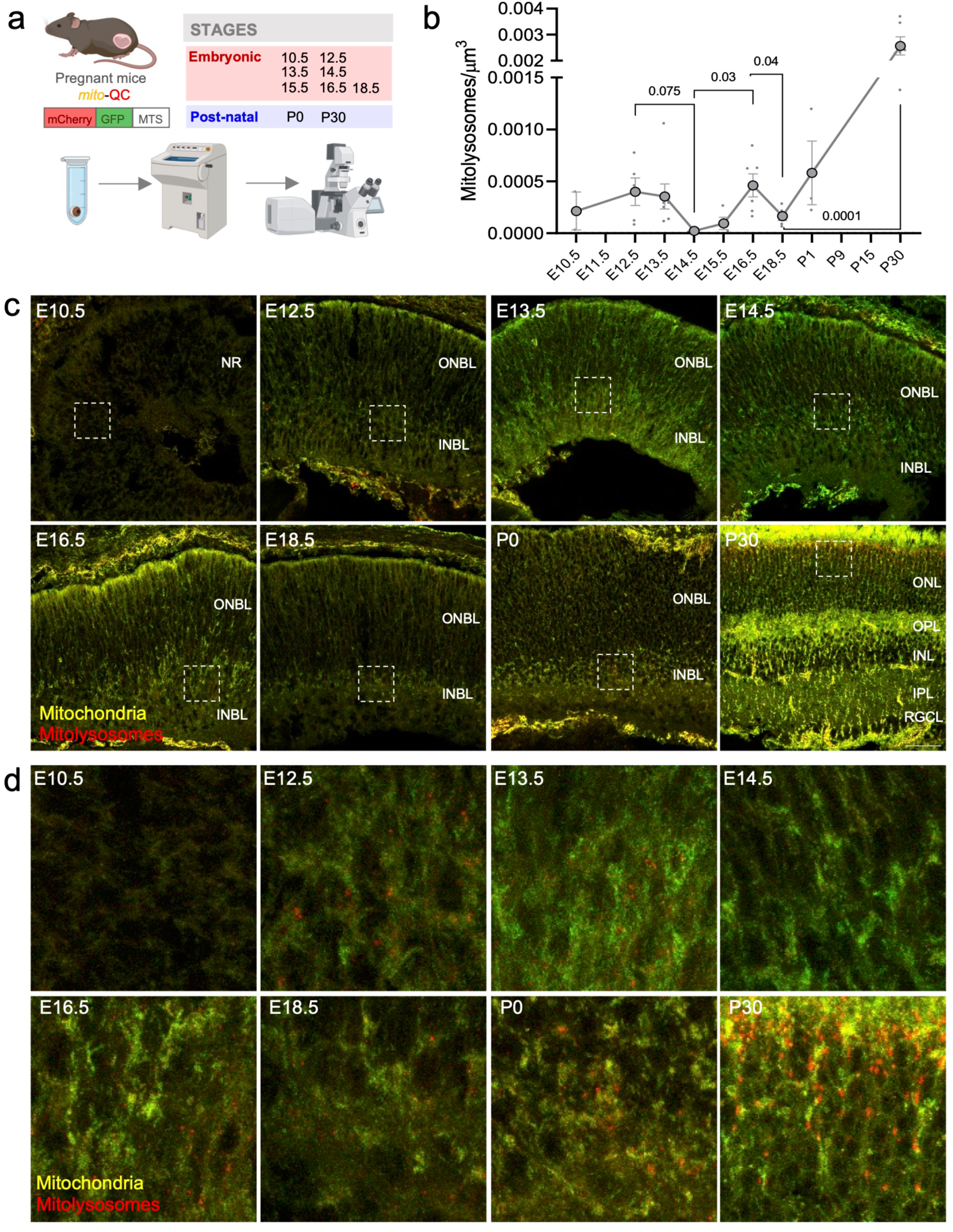
Mitophagy occurs as two distinct peaks during embryonic development, and at higher levels in the mature retina. **(a)** Experimental design. Eyes from mito-QC embryos or pups were collected at each stage and processed for cryostat sectioning. Confocal imaging was performed on all samples simultaneously. **(b)** Number of mitolysosomes per μm^3^ in the neural retina in cryosections from (a). Large dots represent the mean, small gray dots represent individual mice (n = 2–6), and bars represent the SEM. **(c)** Representative images of central retinal cryosections from each developmental stage of mito-QC mice. **(d)** Magnified views of images in (c). Data are presented as mean ± SEM. *P* values were calculated using a two-tailed Student’s *t*-test. Scale bar: 40 μm. E, embryonic day; NR, neural retina; INBL, inner neuroblast layer; INL, inner nuclear layer; IPL, inner plexiform layer; ONBL, outer neuroblast layer; ONL, outer nuclear layer; OPL, outer plexiform layer; P, postnatal day; RGCL, retinal ganglion cells layer; SEM, standard error of the mean.

### PINK1/PRKN-mediated mitophagy is active during retinal development

During the first mitophagy peak, the retina primarily consists of RPCs and RGCs, the first cell type to differentiate [23]. We extracted retinas from E12.5 and E13.5 *mito*-QC embryos, mounted them on nitrocellulose membranes, and captured images in different planes, from the RGC layer to the neuroblast (Nb) layer containing RPCs (**Fig. S4a**). Mitolysosomes were more prevalent in the RGC layer at both stages (**Fig. S4b-d**), supporting a key role of mitophagy in RGCs maturation [7, 24].

Currently, no methods exist for direct measurement of NIX/BNIP3-dependent mitophagy. However, PINK1/PRKN-dependent mitophagy can be assessed by measuring ubiquitin phosphorylation at serine 65 (pUb^Ser65^) [8, 25]. We observed a peak of pUb^Ser65^ at E15.5 and E16.5 (**Fig. 5a, c**), suggesting that the E16.5 mitophagy peak is PINK1/PRKN-dependent, while the first embryonic peak (E12.5–E13.5) and the peak occurring at P30 (i.e. mature retinal mitophagy) are PINK1-independent. PINK1/PRKN pathway activation is typically associated with stress conditions, in order to eliminate damaged mitochondria [8]. Accordingly, we detected a peak in the oxidative stress marker 4-hydroxy-2-nonenal (4-HNE) just before the burst of PINK1/PRKN activity (**Fig. 5b, d**), suggesting that the second embryonic mitophagy peak may be triggered by increased oxidative damage at E14.5–E15.5. To determine whether these mitophagy dynamics are conserved across species, we used a mitophagy reporter zebrafish line (with zebrafish Cox8 fused to EGFP-mCherry) [26]. Two mitophagy peaks were also observed during zebrafish retinal development, at 1-day post-fertilization (dpf) and at 4 dpf (**Fig. 6a, d**). Increased pUb^Ser65^ (Fig. 6b, e) and 4-HNE (Fig. 6c, f) levels at 3 dpf, preceding the second mitophagy peak, suggest involvement of the PINK1/PRKN pathway in zebrafish retinal development.

**Fig. 5.**
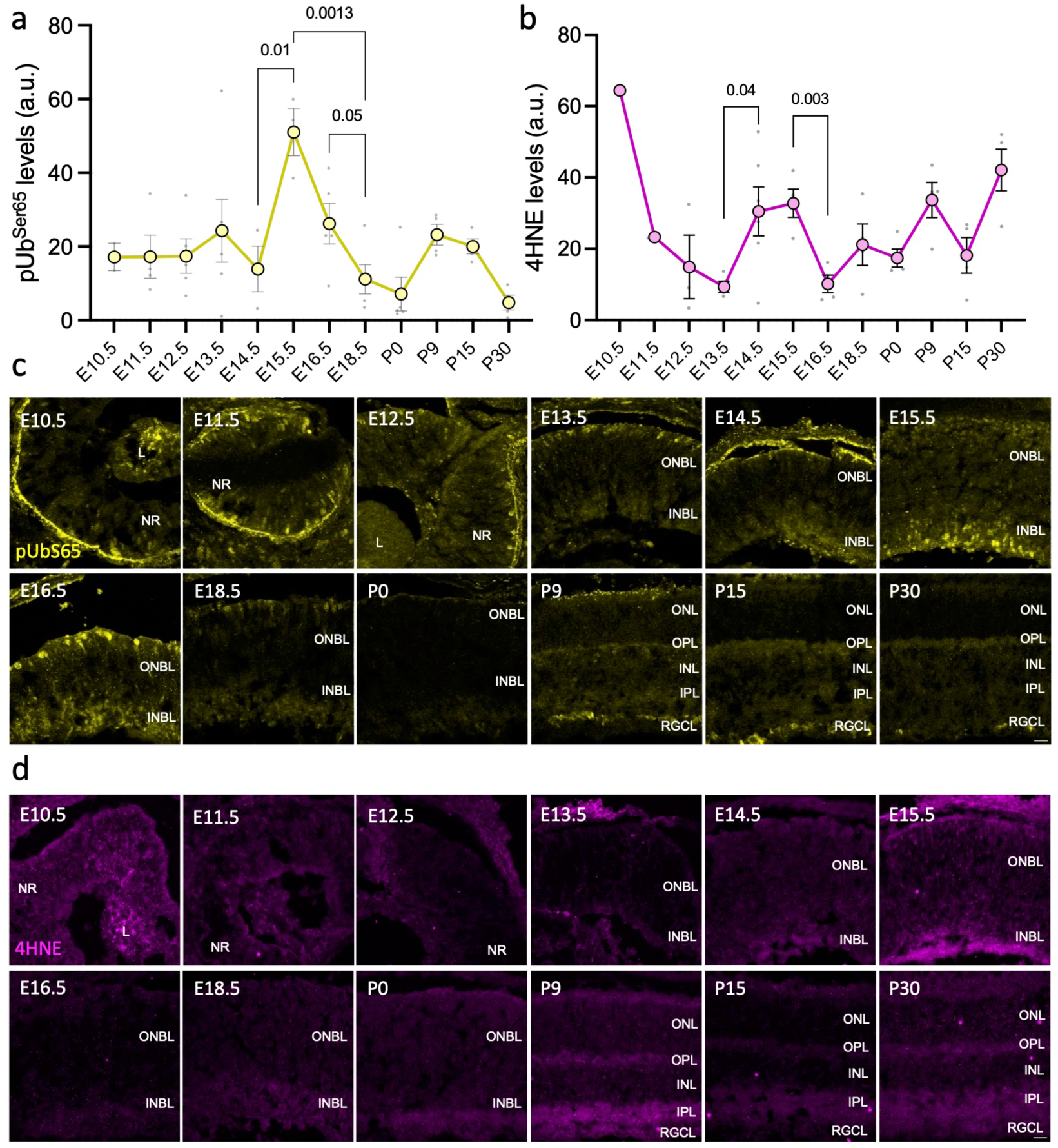
pUb^S65^ and 4-HNE levels are elevated at E15.5, preceding the second mitophagy peak. **(a, b)** Quantification of pUb^S65^ (a) and 4-HNE (b)-positive area in retinal cryosections from different developmental stages, immunolabeled for pUb^S65^ (yellow) and 4-HNE (purple). Large dots represent the mean, and small dots represent individual mice (n = 1–6). **(c, d)** Representative images of central retinal cryosections immunolabeled for pUb^S65^ (c) and 4-HNE (d) at each stage. Data are presented as mean ± SEM. *P* values were calculated using a two-tailed Student’s *t*-test. Scale bar: 20 μm. 4-HNE, 4-hydroxy-2-nonenal; L, lens; NR, neural retina; INBL, inner neuroblast layer; INL, inner nuclear layer; IPL, inner plexiform layer; ONBL, outer neuroblast layer; ONL, outer nuclear layer; OPL, outer plexiform layer; P, postnatal day; pUb^S65^, phosphorylation of ubiquitin at Ser65; RGCL, retinal ganglion cells layer; SEM, standard error of the mean.

**Fig. 6.**
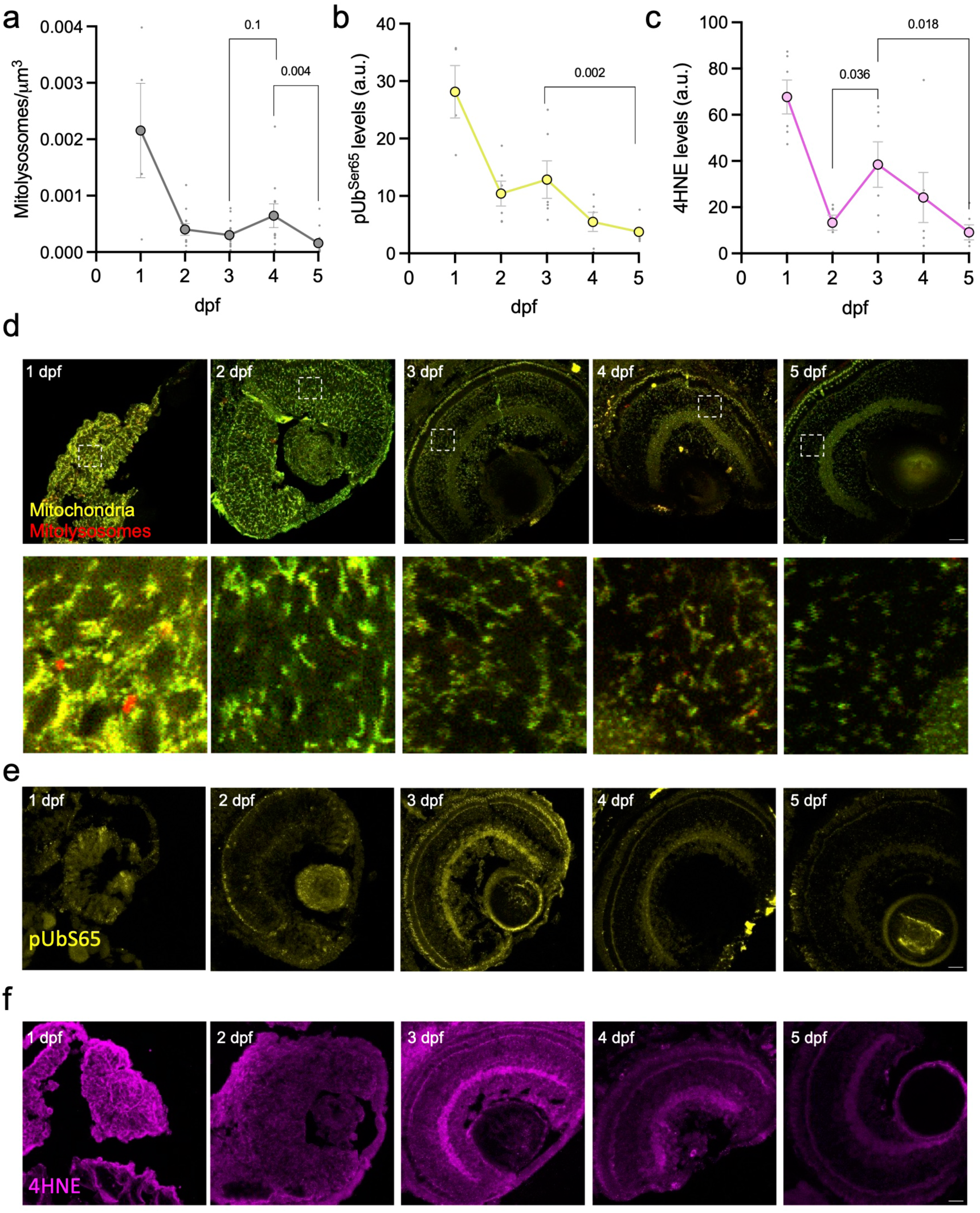
pUb^S65^ and 4-HNE exhibit similar expression patterns during zebrafish eye development. **(a)** Number of mitolysosomes per μm^3^ in the neural retina of cryosections from transgenic tandem-tagged mitofish larvae (Cox8-mCherry-EGFP) at different developmental stages. **(b, c)** Quantification of pUb^S65^ (b) and 4-HNE (c)-positive area in immunolabeled retinal cryosections from distinct developmental stages in zebrafish. Large dots represent the mean, and small dots represent individual fish (n = 3–6). **(d)** Representative images of central retinal cryosections from transgenic tandem-tagged mitofish larvae and magnified views showing mitolysosomes. **(e, f)** Representative images of central retinal cryosections from zebrafish larvae immunolabeled for pUb^S65^ (e, yellow) and 4-HNE (f, purple) at each developmental stage. Data are presented as mean ± SEM. *P* values were calculated using a two-tailed Student’s *t*-test (c) or a two-tailed Mann–Whitney *U*-test (a, b). Scale bar: 20 μm. Dpf, days post fertilization; 4-HNE, 4-hydroxy-2-nonenal; pUb^S65^, phosphorylation of ubiquitin at Ser65; RGCL, retinal ganglion cells layer; SEM, standard error of the mean.

Collectively, these data confirm the importance of mitophagy and autophagy in both mature and developing retinas and support the existence of conserved, stress-induced mitophagy within the physiological context of retinal differentiation.

## Discussion

Here, we present the first comprehensive proteomic analysis of retinal development. Our analysis clearly distinguishes two phases: an embryonic phase characterized by high levels of DNA repair, cell division, and spliceosome-related proteins, and a postnatal stage enriched for proteins involved in synapse formation and metabolism. The marked transition between P0 and P9 likely reflects several critical processes occurring during this period, including retinal astrocyte invasion, retinal vascularization (which reduces hypoxic stress) [27], differentiation of rods, bipolar cells, and Müller glia [28], and synaptogenesis between neuronal layers [5]. Expression of autophagy and mitophagy proteins increased after birth, suggesting a crucial role for these processes in the mature retina. We observed a higher frequency of mitolysosomes in the postnatal retina compared to embryonic time points. However, two small bursts of mitophagy were observed during embryonic development: one at E12.5–E13.5, primarily localized to RGC axons, and another at E16.5 that was preceded by increased oxidative stress and ubiquitin phosphorylation at serine 65 (pUb^Ser65^). These data suggest that PINK1/PRKN-dependent mitophagy may occur in this physiological context as a response to oxidative stress during development. This pattern of oxidative stress followed by PINK1/PRKN-dependent mitophagy was also observed in developing zebrafish retinas, suggesting conservation of this phenomenon across species.

Alternative splicing is strongly associated with differentiation and reprogramming processes [16], and is also observed during early stomach development [29]. This dynamic appears to be conserved in the retina. Autophagy, and specifically mitophagy, are essential for the differentiation of various cell types, including erythrocytes [30, 31], RGCs [7], myoblasts [32], and the transition from beige to white adipocytes [33]. However, we found that autophagy and mitophagy were particularly enriched in the mature retina, supporting housekeeping roles in retinal homeostasis and function, beyond their involvement in differentiation. This is further corroborated by findings in autophagy- and mitophagy-deficient mice, which exhibit alterations ranging from aberrant retinal development to increased susceptibility to external stressors and retinal diseases [6].

Our detailed analysis of mitophagy during retinogenesis using *mito*-QC mice confirmed higher mitophagy levels in the mature retina and the presence of two mitophagy peaks during development. Determining the specific mitophagy pathways involved is challenging and cannot be inferred based on the relative levels of different mitophagy proteins. Therefore, we used pUb^Ser65^ signaling to distinguish between PINK1/PRKN-dependent and -independent mitophagy. Our data indicate that mitophagy in the mature retina at P30 is PINK1/PRKN-independent (**Fig. 5a, c**), consistent with a previous study where it is shown how PINK1 KO retinas display the same mitolysosomes number than WT retinas [34]. This, along with the recent finding that pUb^Ser65^ is only present in aged retinas [25], suggests that PINK1/PRKN mitophagy in the retina may constitute a secondary pathway that is activated in conditions of stress. A similar reasoning can be applied to the embryonic mitophagy peaks. The first peak, occurring at E12.5–E13.5 without pUb^Ser65^ signaling (**Fig. 4b-d**), coincides with the onset of RGC differentiation in the embryonic retina [3] and is largely restricted to RGC axons (**Fig. S4d**). In addition to its role in RGC differentiation [7], we hypothesize that this mitophagy peak may also contribute to RGC axon growth, given that the optic nerve fibers (composed of RGC axons) reach the visual cortex at E16.5. We cannot rule out the involvement of other mitophagy effectors, such as NIX, BNIP3, FUNDC1, or AMBRA1, at these stages. The second mitophagy peak, at E16.5 (**Fig. 4b-d**), is associated with pUb^Ser65^ signaling, which is elevated at E15.5 and E16.5 (**Fig. 5a, c**). Furthermore, the slight downregulation of the PINK1/PRKN negative regulator USP30 at E15.5 (**Fig. 3d**) further supports a role of PINK1/PRKN activation during this mitophagy burst. Oxidative stress plays a critical role in development, influencing cellular signaling, differentiation, and survival [35]. Indeed, retinal 4-HNE levels peak just before this second mitophagy burst (**Fig. 5b, d**). While the involvement of PINK1 in removing the endoplasmic reticulum (ER) and mitochondria has been described during *Drosophila* intestinal development [36], this study provides the first evidence of PINK1/PRKN-mediated mitophagy associated with developmental processes and oxidative stress in mice and zebrafish, previously attributed solely to NIX/BNIP3-dependent mitophagy [7, 30, 37].

## Materials and Methods

### Animal procedures

All animal procedures conformed to the European Union regulations and adhered to the ARVO Statement for the Use of Animals in Ophthalmic and Vision Research. The study was approved by the CSIC ethical review board and the Comunidad de Madrid (PROEX 154.3/21). Mice were housed in the CIB animal facility under a 12-hour light/dark cycle in a temperature-controlled barrier room, with *ad libitum* access to food and water. Both male and female mice were used without distinction. Retinas were consistently collected at 9:00 AM, after lights on, to minimize circadian rhythm-related variations in mitophagy. Timed pregnancies were achieved by single-night mating. *Mito*-QC mice (constitutive knock-in of mCherry-GFP-mtFIS1^101–152^) generated in the laboratory of Ian Ganley [38] were bred in the CIB animal facility. Briefly, recombination-mediated cassette exchange was used to insert a CAG promoter-driven mCherry-GFP-mtFIS1^101–152^ open reading frame, including a Kozak sequence, into the *Rosa26* locus. Mice were maintained on a C57BL/6J background. Transgenic zebrafish were generated as previously described [9].

### Tissue extraction and preparation

For stages E12.5 to P30, right eyes were collected for histological analysis, and left retinas were dissected under a microscope for proteomic analysis. For E10.5 and E11.5, whole heads were collected for histology. Right eyes (or whole heads) were fixed for 1 hour in 3.7% paraformaldehyde (PFA) (171010; EMS) in 200 mM HEPES (4-(2-hydroxyethyl)-1-piperazineethanesulfonic acid), pH 7.0 (15630-080; Gibco), and washed with phosphate-buffered saline (PBS). Left retinas were stored directly at -80°C. Whole eyes were stored in PBS containing 0.01% sodium azide at 4°C until further processing. Zebrafish larvae were fixed overnight in 4% PFA (pH 7.2). Fixed eyeballs or larvae were cryoprotected in a sucrose gradient (15% and 30% in PBS with 0.01% sodium azide) and embedded in optimal cutting temperature (OCT) compound (Tissue-Tek, Sakura Finetek, 4583). Retinal sections (12 μm thick) were cut using a cryostat (Leica Microsystems). For embryonic retinal flat mounts, eyes were enucleated, and retinas were dissected in PBS. Neural retinas were mounted on nitrocellulose membranes with 0.44-μm pores (Sartorius, 11301-80), with the ganglion cell layer facing upward, and carefully adhered to the membrane using forceps. Retinas were fixed for 1 hour in 3.7% PFA (171010; EMS) in HEPES, pH 7.0 (15630-080; Gibco), and washed with PBS.

### Mitophagy assessment and immunofluorescence assays

Mitophagy was assessed in retinal sections and flat mounts. Nuclei were visualized with DAPI (4′,6-diamidino-2-phenylindole, 1 μg/ml; D9542, Sigma), and samples were mounted with Vectashield (H-1000-10, Palex Medical). For immunofluorescence, retinas were permeabilized with 0.15% Triton X-100 (T9284, Sigma) for 30 minutes and blocked for 1 hour with blocking buffer (BGT: 3 mg/ml bovine serum albumin [BSA], 0.25% Triton X-100, 100 mM glycine in PBS). Retinal samples were incubated overnight at 4°C with primary antibodies against OTX2 (1:100; AF1979-SP, R&D Systems), MTCO1 (1:100; 459600, Invitrogen), pUb^Ser65^ (1:100; ABS1513-I, Merck), and 4-HNE (1:100; ab46545, Abcam). After washing with PBS, samples were incubated for 1 hour at room temperature (RT) in darkness with secondary antibodies (1:200; Alexa Fluor 647; Invitrogen, A11075) and DAPI.

### Image analysis and data quantification

All confocal images for each experiment were acquired using consistent laser intensity and photomultiplier settings to minimize variability. DAPI nuclear staining was used solely for field selection during imaging. Images were acquired using a Leica TCS SP8 STED 3X system and processed using Fiji/ImageJ. Experimental design schemes were created using BioRender. Unless otherwise specified, representative images are maximum intensity projections of *z*-stacks, with a *z*-step of 1 μm for whole retinas and cryosections. The percentage area occupied by markers was measured by applying a threshold to the maximum intensity projection of all *z*-stacks. Mitolysosomes were quantified using an ImageJ plugin applying a minimum voxel size threshold [39].

### Protein extraction and proteomic analysis by LC-MS/MS

Proteins were extracted using 1% sodium deoxycholate, reduced with dithiothreitol (DTT) (1 mM final concentration, 30 minutes at RT), and alkylated with iodoacetamide (IAA) (5.5 mM final concentration, 30 minutes at RT in darkness). Samples were digested overnight with trypsin (protease:protein ratio, 1:50). Following digestion, samples were acidified with 50% trifluoroacetic acid (TFA) (approximately 0.5% final concentration, pH < 2) and centrifuged at 4000 rpm for 10 minutes to remove precipitates. Peptides were desalted and purified using StageTips. LC-MS/MS analysis was performed on an Orbitrap Exploris 480 mass spectrometer coupled to a Vanquish NEO UHPLC system (Thermo Scientific). Peptides were separated on a fused silica HPLC column tip (75 μm inner diameter, New Objective, self-packed with ReproSil-Pur 120 C18-AQ, 1.9 μm [Dr. Maisch] to a length of 20 cm) using a gradient of solvent A (0.1% formic acid in water) and solvent B (0.1% formic acid in 80% acetonitrile in water). Samples were loaded with 0% B at a flow rate of 600 nl/min. Peptides were separated using a 5–30% B gradient over 85 minutes at a flow rate of 250 nl/min. Spray voltage was set to 2.3 kV, and ion transfer tube temperature was 250°C. No sheath or auxiliary gas was used. The mass spectrometer was operated in data-independent acquisition (DIA) mode. After each survey scan (mass range *m/z* = 400–1200; resolution: 120,000), 34 DIA scans with an isolation width of 24 *m/z* were performed, covering a precursor range of 400–1200 *m/z*. Automatic gain control (AGC) target value was set to 300%, resolution to 30,000, and normalized stepped collision energy to 25.5%, 27%, and 30%. Raw MS files were analyzed using Spectronaut software (version 19) with the directDIA+ workflow and standard settings, using a UniProt full-length mouse database and common contaminants (keratins, digestive enzymes) as reference. Statistical analysis and graphs were generated using R (v4.2.0) with the following packages: tidyverse, dplyr, ggplot2, pheatmap, FactoMineR, factoextra, matrixTests, patchwork, and ggfortify. *P*-values were adjusted for multiple hypothesis testing using the Benjamini-Hochberg method. Over-representation analysis (ORA) of *k*-means clustered proteins was performed using g:Profiler [40].

### Statistical analysis

Data are presented as mean ± SEM. Statistical analyses were performed using GraphPad Prism software (GraphPad Software, Inc.). Two-tailed Student’s *t*-tests were used for comparisons between treatments or developmental stages. If normality or homoscedasticity assumptions were violated, non-parametric tests (Mann-Whitney *U*-test) were used for two-group comparisons. A significance threshold of *p* < 0.05 (two-tailed) was applied for all tests. Sample sizes are indicated in the respective figure legends.

### Accession number

MS proteomics data have been deposited in the ProteomeXchange Consortium via the PRIDE [41] partner repository with the dataset identifier PXD058997.

Supplementary information is available on the Journal of Molecular Biology website.

## Acknowledgements

We thank Angélica Horrillo and the technical staff at the animal facility, M. Teresa Seisdedos and Gema Elvira at the Confocal Microscopy facility at the CIB Margarita Salas, CSIC. We thank Owen Howard for English editing and all members of the Autophagy Lab and Estela Area’s Lab for thoughtful discussions and support.

## Author Contributions

Conceptualization: J.Z.M., J.I.J.L., and P.B.; data curation: J.Z.M., J.I.J.L. and M.S.; formal analysis: J.Z.M. and J.I.J.L.; funding acquisition: P.B.; investigation and methodology: J.Z.M., J.I.J.L., M.S., B.V.Z., S.A.G., P.T., B.J.M.; project administration: P.B.; supervision: I.G., J.D., A.S. and P.B.; validation: P.B.; writing – original draft: J.Z.M.; writing – review and editing: J.Z.M., J.I.J.L., M.S., B.V.Z., B.J.M., I.G., A.S., J.D., and P.B. All authors read and approved the final version of the manuscript.

## Ethics approval

Our studies did not include human participants, human data, or human tissue. All-animal related studies and procedures were performed following European Union guidelines and the ARVO Statement for the Use of Animals in Ophthalmic and Vision Research. Animal procedures were approved by the CSIC ethical review committee and the Comunidad de Madrid (PROEX 154.3/21).

---

**Fig. S1.**
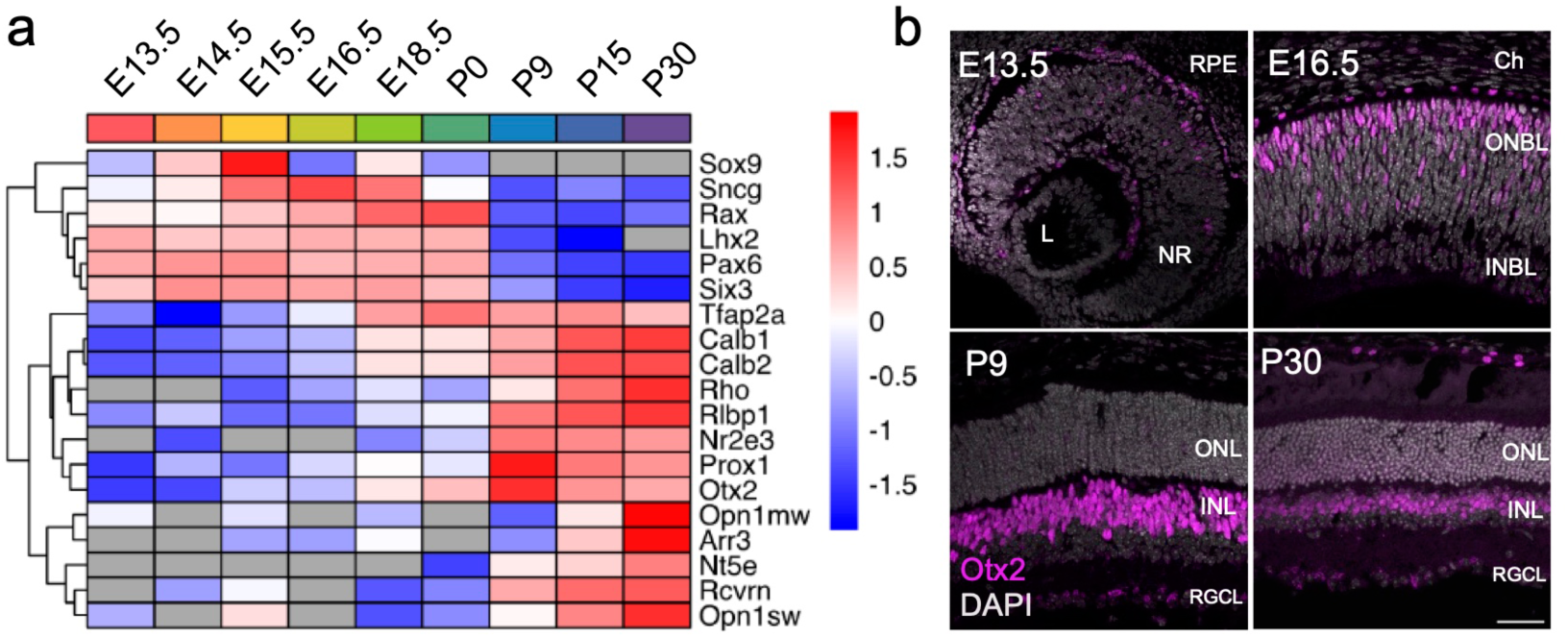
Cell differentiation markers display expected patterns during retinal development. **(a)** z-score hierarchical clustering heatmap visualization of differentiation and cell-specific markers during retinal development. The color scale represents protein expression levels, ranging from dark red (high expression) to dark blue (low expression). **(b)** Representative confocal images of central retinal cryosections immunolabeled for OTX2 (purple) and counterstained with DAPI (gray) at different retinal stages. Ch, choroids; E, embryonic day; L, lens; NR, neural retina; INBL, inner neuroblast layer; INL, inner nuclear layer; ONBL, outer neuroblast layer; ONL, outer nuclear layer; OTX2, orthodenticle homeobox 2; P, postnatal day; RGCL, retinal ganglion cells layer; RPE, retinal pigment epithelium.

**Fig. S2.**
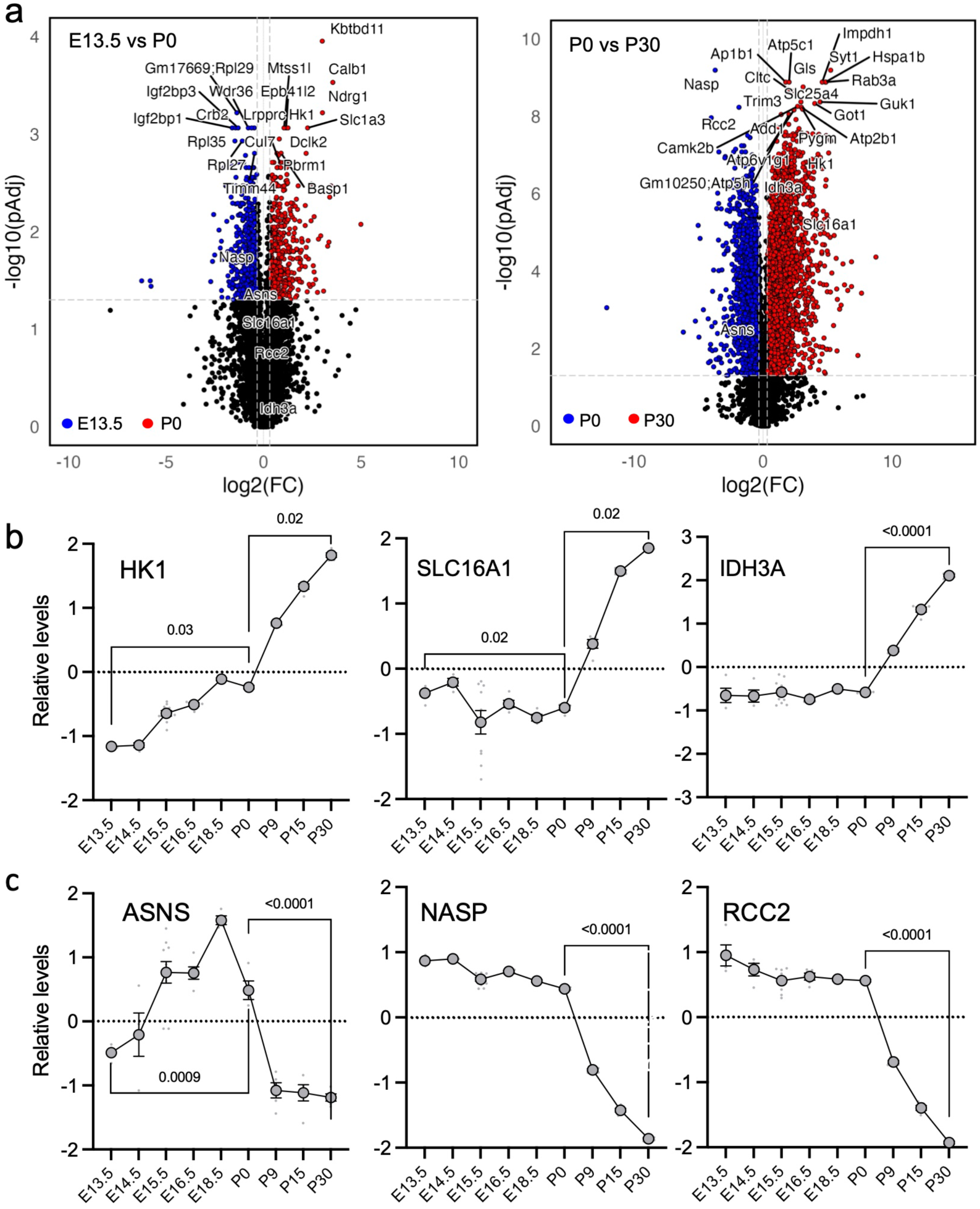
Top hits in volcano plot analysis during embryonic and postnatal retinal development. **(a)** Volcano plots showing the main changes during the first and second development stages. Genes significantly enriched in each stage are highlighted in color as indicated in the legend, based on a *p*-value threshold of <0.05 and a log_2_ fold change cutoff of ±1. Non-significant protein changes are shown in black. **(b)** Z-score normalized expression levels of the metabolic proteins HK1, SLC16A1, and IDH3A as detected by proteomics. **(c)** Z-score normalized expression levels of the DNA replication and repair-related proteins ASNS, NASP, and RCC2 as detected by proteomics. *P* values were calculated using a two-tailed Student’s *t*-test (ASNS, IDH3A, NASP, and RCC2) or a two-tailed Mann–Whitney *U*-test (HK1 and SLC16A1). FC, fold change.

**Fig. S3.**
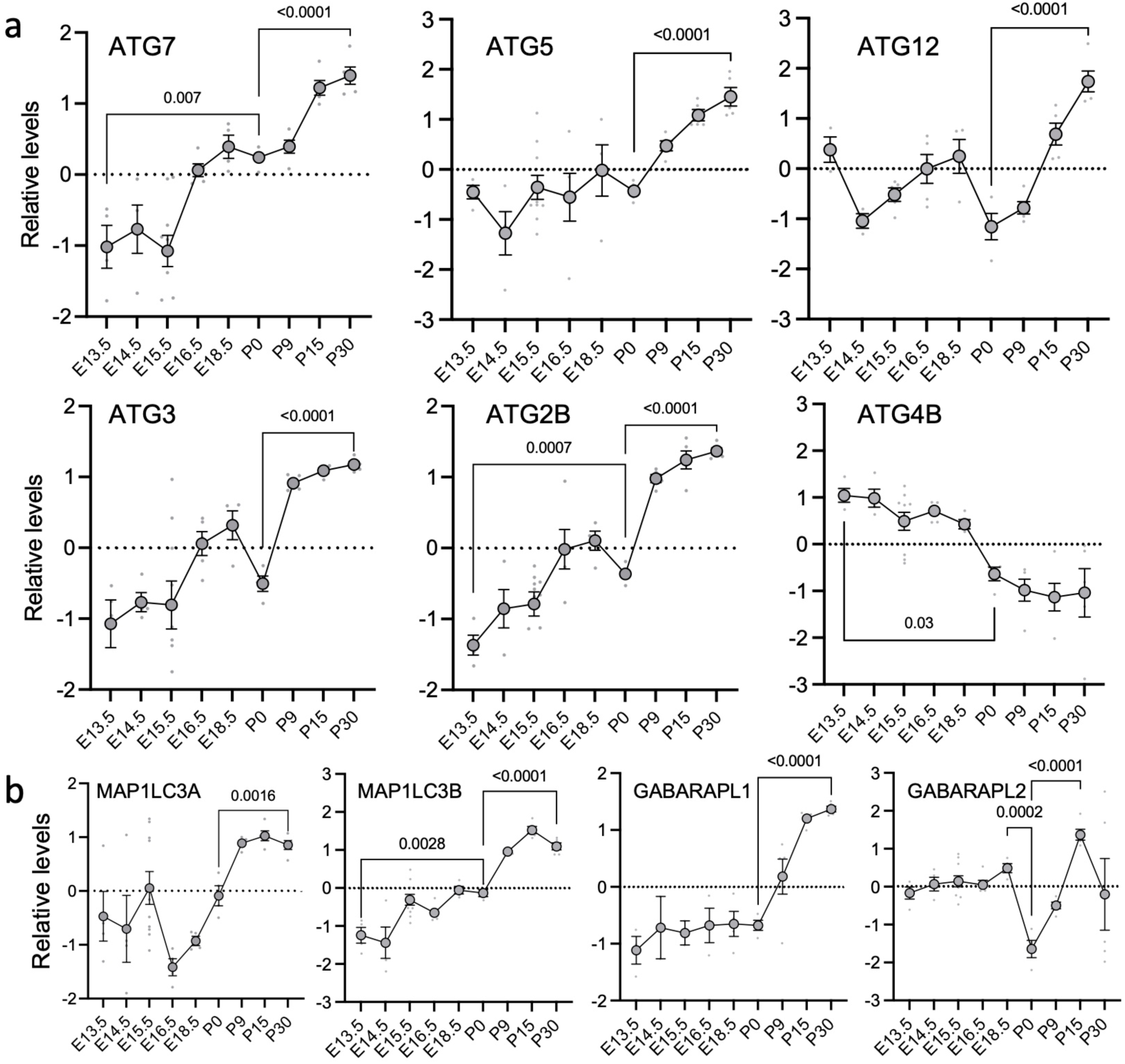
Autophagy machinery proteins are upregulated during retinal development. (**a, b**) Z-score normalized expression levels of the lipidation machinery proteins (a) and mammalian ATG8 proteins (b) detected by proteomics. Large dots represent the mean, small gray dots represent individual mice (n ≥ 4), and bars represent the SEM. *P* values were calculated using a two-tailed Student’s t-test (ATG7, ATG5, ATG12, ATG3, ATG2B, and b) or a two-tailed Mann–Whitney *U*-test (ATG4B).

**Fig. S4.**
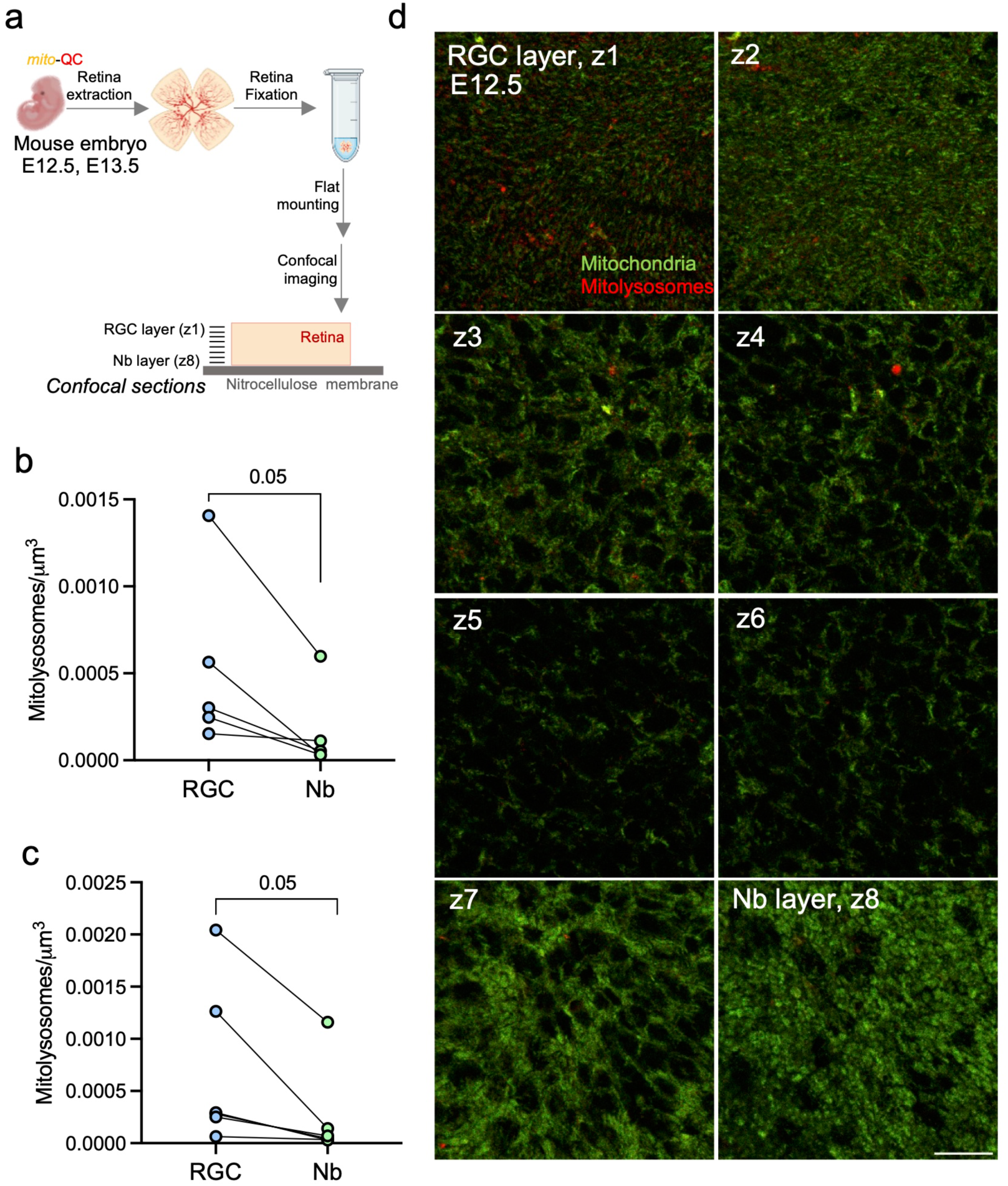
Mitolysosomes are localized to the RGC layer at E12.5 and E13.5. (**a**) Experimental design. Retinas were extracted from mito-QC E12.5 and E13.5 embryos, fixed, and mounted on nitrocellulose membranes before confocal imaging. (**b, c**) Number of mitolysosomes per μm^3^ in the RGC and Nb layers from E12.5 (b) and E13.5 (c) retinas (n = 5–6). (**d**) Representative confocal images taken from the RGC layer towards the Nb layer from an E12.5 mito-QC retina. Data are expressed as the mean ± SEM. P values were calculated using a two-tailed paired Student’s t-test. Scale bar: 20 μm. Nb, neuroblast; RGC, retinal ganglion cell.

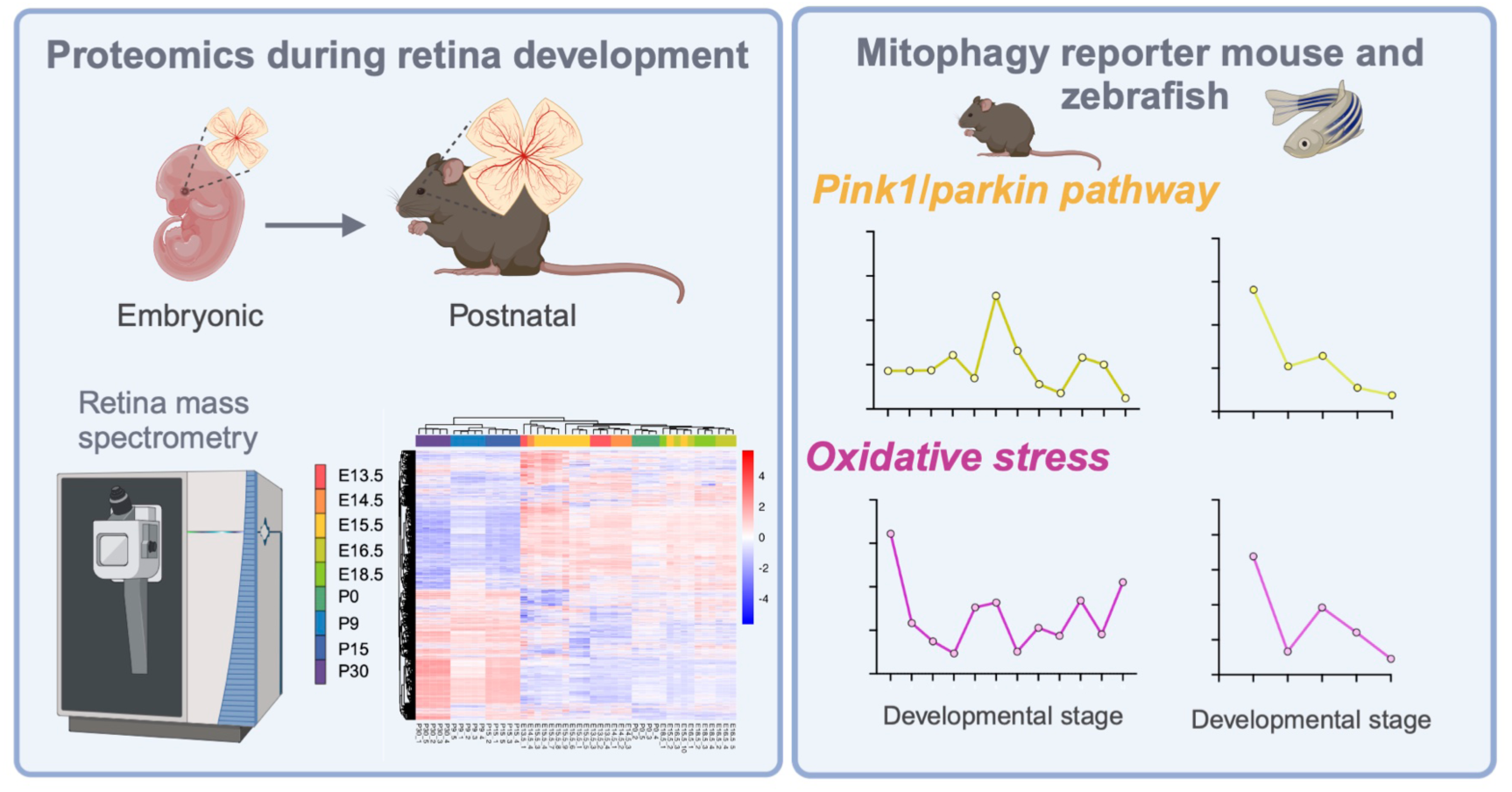

